# ATTRv-V30M Type A amyloid fibrils from heart and nerves exhibit structural homogeneity

**DOI:** 10.1101/2024.05.14.594028

**Authors:** Binh An Nguyen, Shumaila Afrin, Anna Yakubovska, Virender Singh, Jaime Vaquer Alicea, Peter Kunach, Preeti Singh, Maja Pekala, Yasmin Ahmed, Maria del Carmen Fernandez-Ramirez, Luis O. Cabrera Hernandez, Rose Pedretti, Parker Bassett, Lanie Wang, Andrew Lemoff, Layla Villalon, Barbara Kluve-Beckerman, Lorena Saelices

## Abstract

ATTR amyloidosis is a systemic disease characterized by the deposition of amyloid fibrils made of transthyretin, a protein integral to transporting retinol and thyroid hormones. Transthyretin is primarily produced by the liver and circulates in blood as a tetramer. The retinal epithelium also secretes transthyretin, which is secreted to the vitreous humor of the eye. Because of mutations or aging, transthyretin can dissociate into amyloidogenic monomers triggering amyloid fibril formation. The deposition of transthyretin amyloid fibrils in the myocardium and peripheral nerves causes cardiomyopathies and neuropathies, respectively. Using cryo-electron microscopy, here we determined the structures of amyloid fibrils extracted from cardiac and nerve tissues of an ATTRv-V30M patient. We found that fibrils from both tissues share a consistent structural conformation, similar to the previously described structure of cardiac fibrils from an individual with the same genotype, but different from the fibril structure obtained from the vitreous humor. Our study hints to a uniform fibrillar architecture across different tissues within the same individual, only when the source of transthyretin is the liver. Moreover, this study provides the first description of ATTR fibrils from the nerves of a patient and enhances our understanding of the role of deposition site and protein production site in shaping the fibril structure in ATTRv-V30M amyloidosis.

## Introduction

ATTR amyloidosis is a form of systemic amyloidosis caused by the misfolding and amyloid deposition of the protein transthyretin^1^. In its native form, transthyretin is responsible for retinol and thyroid hormone transportation. Transthyretin is predominantly synthesized in the liver, where it enters the bloodstream. It is also synthesized by the retinal epithelium and secreted into the vitreous humor in the eye. Furthermore, it is synthesized in the choroid plexus, where it contributes to the makeup of cerebrospinal fluid ^2^. Although transthyretin circulates as a tetramer, factors like mutations or aging can lead to its conversion to amyloidogenic transthyretin (ATTR), prone to dissociating into monomers that ultimately self-assemble to form amyloid fibrils^3^. These amyloid deposits can be found in the myocardium and nerves, resulting in cardiomyopathies and neuropathies, respectively^4^. In wild-type ATTR (ATTRwt) amyloidosis, the aggregation of wild-type transthyretin leads primarily to cardiac deposition and cardiomyopathy^5^. In variant ATTR (ATTRv) amyloidosis, over 140-point mutations drive aggregation in multiple organs, exhibiting a broad phenotypic variation that is poorly understood^6^.

Structural insights into amyloid fibrils have been instrumental in understanding disease classification in neurodegenerative disorders^7^ and may offer significant insights into the link between structure and phenotype in ATTR amyloidosis. Much of the current understanding of the atomic structure of ATTR fibrils stems from studies on *ex-vivo* extracts^8–14^. These studies reveal that all ATTR fibrils to date share a common conformation made of two transthyretin fragments, with additional local structural variations^10^ and/or multiple protofilaments^8,14^ that could be influenced by the location of the transthyretin mutation. The local variations are mainly found in the residues Gly 57 to Gly 67 (previously defined as *gate*) that close, open, or block a polar cavity formed by residues Leu 58 to Ile 84^10^. Variations in the gate have been found in ATTR-I84S cardiac fibrils^10^ and ATTRv-V30M fibrils extracted from vitreous humor^8^. ATTR fibrils with multiple protofilaments have been found in the same ATTRv-V30M vitreous humor fibrils as well as in ATTR-V122Δ cardiac fibrils^8,14^. The structures of ATTRwt fibrils and other ATTRv fibrils display a single protofilament and a closed gate, and are structurally consistent ^9,11–13^.

The variation found in ATTRv-V30M fibrils extracted from multiple organs (heart vs. vitreous humor) of two independent patients^8,11^ may suggest that fibrils polymorphism may be driven by the tissue environment. Building upon this observation, in another recent study, we investigated the structures of fibrils extracted from multiple organs from ATTRv-T60A amyloidosis patients and found a structural consistency across the thyroid, heart, liver, and kidney^15^. These discoveries raised the following questions. Does ATTRv-V30M polymorphism depend on the source of transthyretin (retinal epithelium vs liver)? Or the individual? To contribute to the understanding of these questions, we set out to determine the structures of ATTR fibrils obtained from the cardiac and nerve tissues of the same ATTRv-V30M amyloidosis patient. These fibrils are produced by the liver but deposited in different organs. Our comparative analysis revealed the same fibril structure in both organs, characterized by a single protofilament and a closed gate. This consistency underscores the uniformity of fibril structures within multiple organs of the same patient, when the precursor protein is produced by the liver. Our study provides the first structure of ATTR fibrils collected from the nerves of a patient.

## Results

### Fibril extraction and cryo-EM data processing

We obtained freshly frozen cardiac and nerve tissue samples from an ATTRv-V30M amyloidosis patient. For fibril extraction, we used a gentle water-based extraction method as previously described ^10,11^. We confirmed the fibril type (type A vs Type B) using an in-house antibody directed against a C-terminal fragment of transthyretin characteristic of type A fibrils ^16^ (Fig. 1a). We also used mass spectrometry to confirm the presence of both wild-type and mutant transthyretin, as expected from patients’ zygosity. We identified several non-tryptic proteolytic sites within both the disordered regions and the amyloid core for both the nerve and cardiac fibrils, although there were more cleavage sites in the cardiac tissue compared to the nerve tissue (Supplementary Figure 1) We then assessed the quality and distribution of fibrils using negative stain electron microscopy (Fig 1b-c). Subsequently, we screened and optimized samples for cryo-EM structure determination (Fig. 1d-e).

**Figure 1:**
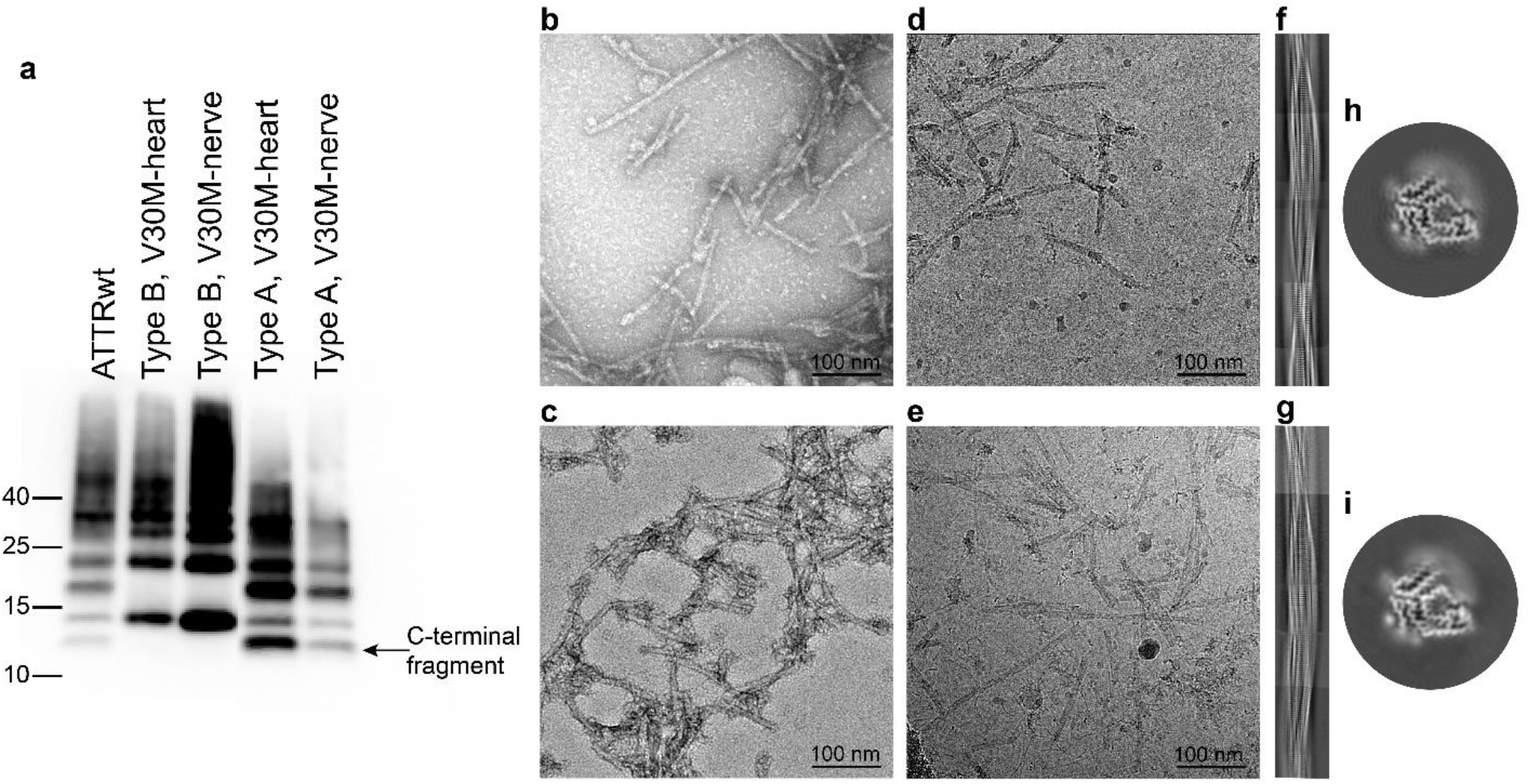
ATTRv-V30M fibrils from heart and nerve. (a) Anti-TTR western blot extracted using an antibody that detects transthyretin C-terminus fragments. This assay is used to confirm fibril type (Type A vs Type B). Type B fibrils were used as control. Negative stain images of fibrils extracted from the (b) nerve and (c) heart tissues. Representative cryo-EM micrographs from the patient (d) nerve (e) and cardiac fibril datasets. Representative 2D class averages depicting the fibril crossover for (f) nerve and (g) heart. 3D class averages of fibrils from (h) nerve and (i) heart.

### ATTRv-V30M fibrils from the heart and nerve of the same patient are structurally similar

We collected cryo-EM images of ATTRv-V30M fibrils extracted from the nerve and cardiac tissue of the ATTRv-V30M amyloidosis patient. From 2D class averaging, we observed two main fibril types that vary in the presence or absence of a twist. The untwisted fibrils represented a minor percentage of the total fibril population, and because of their lack of a twist, were unsuitable for helical structure reconstruction (Supplementary table 1). The dominant species had a discernible twist and were used for 2D class stitching (Figure 1f-g), 3D classification (Figure 1h-i), and helical structure reconstruction (Figure 2a-f).

**Figure 2:**
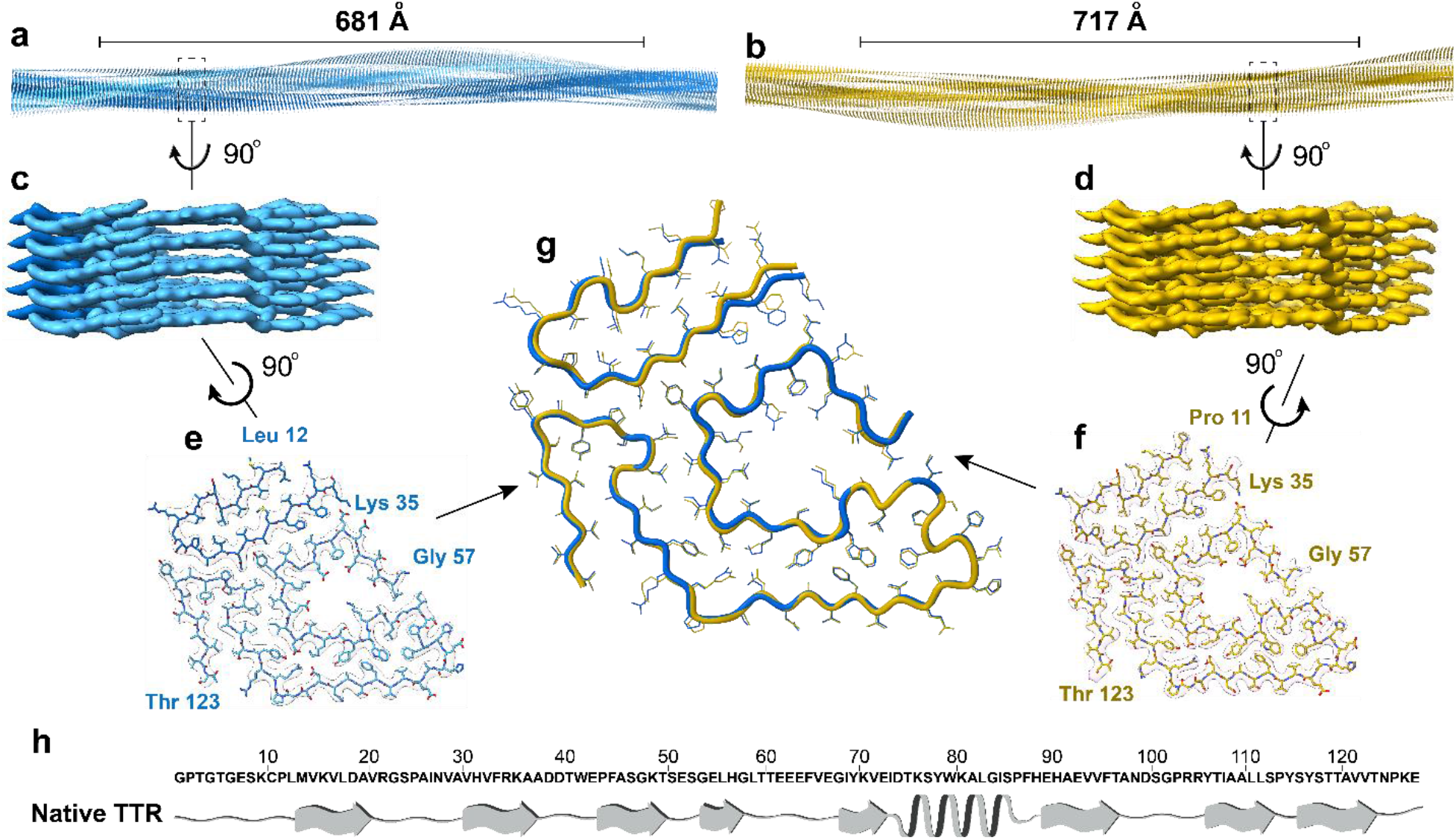
Cryo-EM structural analysis of ATTRv-V30M fibrils from heart and nerve. Cryo-EM model obtained from helical reconstruction, depicting the full crossover in nerve (a) and (b) heart. An enlarged section showing the side view of the density map for the (c) nerve and (d) heart. Cryo-EM density and atomic model displaying the structure of the fibril from (e) nerve and (f) heart, encompassing two fragments of transthyretin, an N-terminus fragment (residues Pro 11 or Leu 12 to Lys 35) and a C-terminus fragment (residues Gly 57 to Thr 123). (g) A top view of the structural backbone alignment of nerve (colored blue) and cardiac (colored yellow) ATTR fibril structure showing an r.m.s.d of 0.621Å. (h) Primary sequence of wild-type transthyretin (at the top), accompanied by the secondary structures of its native-folded form (PDB 4TLT).

3D classification of type A ATTRv-V30M fibrils from the nerve revealed the presence of a single class of fibrils (Fig 1h). Upon refining, this class achieved a resolution of 3.2Å as estimated by FSC curves (Supplementary Fig. 2). Comprising a single protofilament, these fibrils exhibit a separation of 4.93Å between layers and a crossover distance of 681 Å (Fig 2a, 2e). This structural configuration closely resembles the published structure of cardiac fibrils from ATTRv-V30M, ATTRwt, and several ATTRv amyloidosis patients, with a closed gate in the polar cavity ^9,11–13^. Briefly, the structure encompasses an N-terminal fragment spanning residues Pro 11 to Lys 35, alongside a C-terminal fragment spanning residues Gly 57 to Thr 123. It encloses a polar channel from residue Gly 57 to Ile 84, with a gate formed by residues Gly 57 to Gly 67 (Figure 2e). The density map does not fit residues Ala 36 to His 56, suggesting that they may be disordered and/or absent. Similarly, we do not observe any unsatisfied density inside the polar channel, which may signify that its content is not ordered and/or the resolution of the density map is not sufficient to discern occupying molecules.

3D classification of type A ATTRv-V30M fibrils from the heart also yielded a single 3D class (Fig 1i). Cryo-EM reconstruction of the 3D map resulted in a resolution of 3.1 Å, as estimated by FSC curves (Supplementary Fig 2). These fibrils also comprised of a single protofilament, with a separation between layers of 4.90 Å and a crossover distance of 717 Å (Fig 2b, 2f). The refined structure of ATTRv-V30M fibrils from the heart resembles the structure of ATTRv-V30M fibrils from the nerves of the same patient. The only observed difference was the lack of residue Pro 11 in the nerve fibrils (Fig 2e-h). In fact, the all-atom root mean square deviation (r.m.s.d.) calculated between the two fibril structures from the type A ATTRv-V30M patient was 0.621 Å (Figure 2g-h).

As expected, solvation energy calculations show that the fibril structures from both nerve and cardiac fibrils exhibit similar stability. The fibril structure from the nerve has free energies per residue and per chain recorded as −0.68 kcal/mol and −62.1 kcal/mol respectively, while the nerve fibrils show values of −0.73 kcal/mol and 67.5 kcal/mol per residue and per chain, respectively (as depicted in Supplementary Figure 3a-b). This level of stability is comparable to that of previously documented ATTRv and ATTRwt fibril structures^9,10,14^.

**Figure 3:**
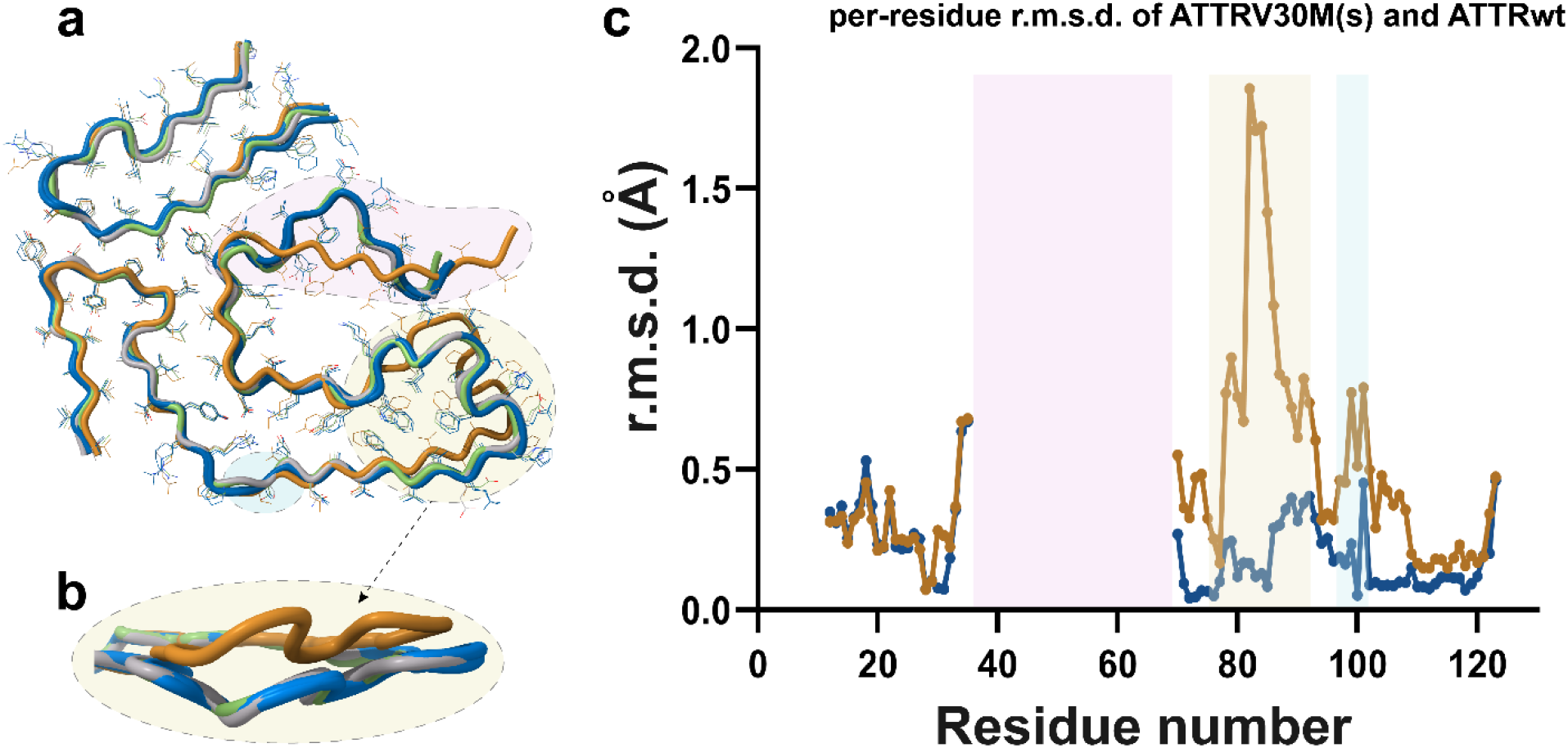
Structural comparison of ATTR fibril structures. A structural backbone alignment of ATTR fibrils is presented with (a) top view (b) zoomed-in side view. The ATTRv-V30M from this study is shown in blue, previously published ATTRv-V30M in green, ATTRwt in grey, and ATTRv-V30M from the eye in orange. A dot plot of the RMSD analysis (comparing only the Cα in Å per residue) featuring two groups: one including ATTRv-V30M from this study, previously published cardiac ATTRv-V30M, and previously published ATTRwt grouped together represented in yellow, and another for vitreous humor ATTRv-V30M depicted in orange (c). Each dot in the plot represents the average r.m.s.d for each group compared to a consensus sequence calculated by GESAMT^17^.

Further analysis of solvation energy while incorporating residue composition information (Supplementary Figure 3c-d) in the nerve and cardiac fibril structure highlights three significant hydrophobic pockets that enhance fibril stability. These include the inner interface formed by a hairpin between residues Leu 12 and Val 32 in the N-terminal fragment, an aromatic pocket around residues Trp 79 and Phe 95 in the C-terminal fragment, and a triquetra linking the C- and N-terminal fragments at the core’s center. The structure’s stability is further enhanced by backbone and side chain interactions, such as hydrogen bonds and π-π stacking, similar to those found in other ATTR fibrils structures^9,10^.

### Structural comparison of ATTRv-V30M fibrils from the nerves, heart and the vitreous humor of the eye

We aligned the structures of all published ATTRwt structures and cardiac, nerve and vitreous humor ATTRv-V30M fibrils including the heart and nerve solved in this study (Figure 3a-b) and measured backbone displacement (limited to the Cα atoms) using root mean square deviation (r.m.s.d) calculations with GESAMT^17^. We excluded residues Leu 58 to Glu 66 in the gate region of the polar channel from our analysis, recognizing their role in influencing ATTR fibril polymorphism, as previously described^10^. In ATTRv-V30M nerve and heart from this study and both the ATTRwt and previously published ATTRv-V30M hearts, the r.m.s.d of the Cα atoms of the fibril structure, show differences less than 0.5 Å, indicating high structural similarity. However, in the model representing the fibril structure from ATTRv-V30M vitreous humor, certain regions exhibit deviations ranging from 0.5 to 2 Å, notably in the backbone region encompassing residues Ala 81 to His 90 (Fig 3b). This marked deviation is attributed to the absence of interaction between Leu 58 and Iso 84, which destabilizes the backbone.

## Discussion

In this work, we describe for the first time the structures of fibrils extracted from nerves of an ATTR amyloidosis patient and compare these with fibril structure from the same patient’s heart. We found that ATTRv-V30M fibrils from these organs share a common conformation. Their structures are characterized by the presence of a single protofilament made of two transthyretin fragments, where the C-terminus forms a polar channel closed by a gate made of residues Gly 57 to Gly 67. Our findings underscore the structural uniformity observed in ATTRv-V30M fibrils from both the heart and nerve tissues of the same patient. Together with other previous observations, these results open new inquiries about the influence of the protein deposition or production site on the structure of ATTRv-V30M fibrils.

Cryo-EM analysis of cardiac fibrils extracted from primarily cardiomyopathic patients reveals a consistent structural landscape^9,13^. Conversely, cardiac fibrils extracted from primarily polyneuropathy cases, such as ATTRv-I84S^10^ or ATTRv-V122Δ^14^, exhibit structural polymorphism both within and between patient sample. A multi-organ study of T60A, a mutation associated with a mixed phenotype with cardiomyopathy and polyneuropathy, shows structural homogeneity in fibrils extracted from various organs of the same patient^15^. This is consistent with our present work on late-onset ATTRv-V30M patients, predominantly cardiomyopathic, where we observe structural homogeneity in fibrils from different organs (Fig. 4). The structural comparison of all ATTR fibrils determined to date hints at a conceivable association between fibril polymorphism and a more neuropathic phenotype. This association needs to be validated with a larger sample size, including multiple variants, tissue samples, and phenotypes.

**Figure 4:**
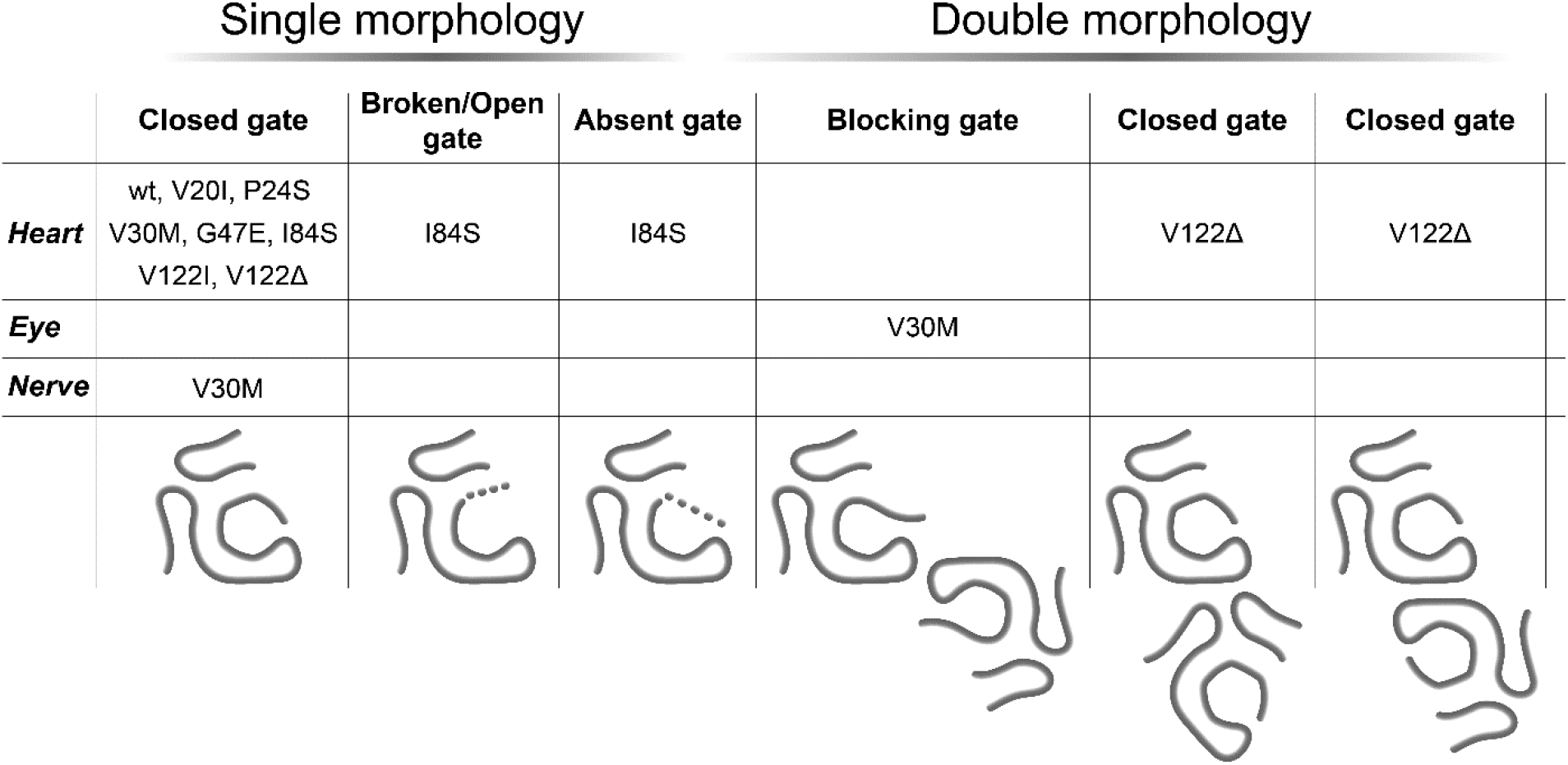
Schematic representation of the structural polymorphism detected in ATTR fibrils across various organs and mutations to date.

Cryo-EM structures of amyloid fibrils have shown that a single protein sequence can acquire multiple amyloid folds^18^. Exemplifying this observation, others previously showed that ATTRv-V30M fibrils deposited in the vitreous humor have the same sequence but different structural conformation than those deposited in the heart^8,11^. These differences could also be driven by several factors. One is the influence of the site of protein production; Transthyretin found in amyloid deposits within the vitreous humor originates from the retinal epithelium, whereas the transthyretin present in cardiac amyloid deposits is synthesized by the liver^19^. A second factor is the influence of the microenvironment on protein aggregation. A previous study shows that the same sequence can lead to different structures depending on the local region of deposition within the same organ. In particular, serum Amyloid A (SAA) amyloid fibrils extracted from the glomerular and vascular regions of the kidney do not share a common structure ^20,21^. To investigate whether the observed structural variations in ATTRv-V30M fibrils are influenced by the site of deposition— and not the production—, we analyzed fibril structures derived from two different tissues of the same patient. In both tissues, the source of the deposited transthyretin is the liver^19^. We found that the structures of fibrils from the two tissues are quasi-identical, suggesting that at least in the case of this ATTRv-V30M patient, the site of deposition may not influence the structure of the fibril. Instead, ATTR fibril structure may be pre-determined in earlier stages of the protein aggregation pathway.

Our findings suggest that the final conformation of systemic ATTR fibrils may be influenced by the inherent properties of the monomer and/or the unique conditions present at the time of the initial misfolding or early aggregation events^22^. The presence of consistent fibril conformation across multiple organs^15^ suggests that the structure of ATTR fibrils is somewhat footprinted in the body of the patient, prior to deposition. The Kelly group and our group have identified nonnative and aggregated species what circulate in the blood of ATTR amyloidosis patients ^23,24^. These species, perhaps, may adopt a predetermined conformation that could serve as a template for further aggregation and amyloid deposition. Further studies exploring this possibility would be beneficial in understanding the complexity of the ATTR amyloidosis phenotype.

## Materials and methods

### Patient samples

Cardiac and peripheral nerve tissues were acquired postmortem from a male ATTR patient with the V30M mutation. Patient was diagnosed with heart failure and neuropathy and died at the age of 60. Control type B samples were obtained from a male ATTR patient who died at the age of 52. All samples were anonymized and provided by Dr. Merrill D. Benson’s laboratory at the University of Indiana.

### Extraction of amyloid fibrils from human cardiac and peripheral nerve tissues

ATTR amyloid fibrils were extracted from fresh frozen human tissue^10,11,25^. Approximately 200 mg of frozen cardiac or peripheral nerve tissues was thawed, cut into fine pieces, and suspended in 1 mL Tris-calcium buffer (20 mM Tris, 150 mM NaCl, 2 mM CaCl_2_, 0.1% NaN_3_, pH 8.0). The mixture was then centrifuged for 5 minutes at 3100 × g and 4°C, and the resulting pellet was washed four times with the same buffer. Subsequently, the pellet was resuspended in 1 mL of 5 mg/mL collagenase solution (collagenase was dissolved in Tris-calcium buffer) and incubated overnight at 37°C, shaking 400 rpm. After another round of centrifugation, the pellet was resuspended in 1 mL of Tris–EDTA buffer (20 mM Tris, 140 mM NaCl, 10 mM EDTA, 0.1% NaN3, pH 8.0) and washed nine more times. Following the final wash, the pellet was resuspended in 200 μL of ice-cold water with 5-10 mM EDTA and centrifuged to collect amyloid fibrils in the supernatant. This extraction step was repeated five times. Material from different organs were handled and analyzed separately.

### Typing and Western blotting of extracted ATTR fibrils

To confirm the fibril type, western blotting was performed on the extracted fibrils. In brief, 0.5 µg of fibrils were dissolved in a tricine SDS sample buffer and boiled for 2 minutes at 85 °C. Subsequently, they were loaded onto a Novex™ 16% tris-tricine gel system using a Tricine SDS running buffer. The determination of TTR type was achieved by transferring the gel contents to a 0.2 µm nitrocellulose membrane, followed by probing with a primary antibody (diluted 1:1000) targeting the C-terminal region of the wild type TTR sequence (GenScript). Horseradish peroxidase-conjugated goat anti-rabbit IgG (Invitrogen, diluted 1:1000), was used as a secondary antibody. TTR content was visualized using Promega Chemiluminescent Substrate, following the manufacturer’s instructions.

### Mass Spectrometry (MS) sample preparation, data acquisition and analysis

For tryptic MS analysis, 0.5 µg of extracted ATTR fibrils were dissolved in a tricine SDS sample buffer, heated for 2 minutes at 85 °C, and separated on a Novex™ 16% tris-tricine gel system using a Tricine SDS running buffer. After staining with Coomassie dye and subsequent destaining, the ATTR smear was excised from the gel for MS analysis. The excised samples were then subjected to overnight digestion with trypsin (Pierce), preceded by reduction and alkylation with DTT and iodoacetamide (Sigma–Aldrich). Subsequently, the samples underwent solid-phase extraction cleanup utilizing an Oasis HLB plate (Waters), after which they were injected onto a Q Exactive HF mass spectrometer coupled to an Ultimate 3000 RSLC-Nano liquid chromatography system. Injection onto a 75 µm i.d., 15-cm long EasySpray column (Thermo) ensued, followed by elution with a gradient from 0% to 28% buffer B over a span of 90 minutes. Buffer A consisted of 2% (v/v) ACN and 0.1% formic acid in water, while buffer B was composed of 80% (v/v) ACN, 10% (v/v) trifluoroethanol, and 0.1% formic acid in water. The mass spectrometer operated in positive ion mode with a source voltage of 2.5 kV and an ion transfer tube temperature of 300 °C. MS scans were conducted at 120,000 resolution in the Orbitrap, with up to 20 MS/MS spectra obtained in the ion trap for each full spectrum acquired using higher-energy collisional dissociation (HCD) for ions with charges 2-8. Dynamic exclusion was set for 20 s after an ion was selected for fragmentation.

The raw MS data files were analyzed using Proteome Discoverer v3.0 SP1 (Thermo). Peptide identification was conducted using a semitryptic search with Sequest HT against the human reviewed protein database from UniProt. Fragment and precursor tolerances of 10 ppm and 0.02 Da were specified, with three missed cleavages allowed. Carbamidomethylation of Cys was designated as a fixed modification, while oxidation of Met was considered a variable modification. The false-discovery rate (FDR) cutoff for all peptides was set at 1%. The mass spectrometry proteomics data have been uploaded to MassIVE (a member of ProteomeXchange) under the accession code MSV000094746.

### Negative-stained transmission electron microscopy

Amyloid fibril extraction was confirmed using transmission electron microscopy. Briefly, a 3 μL sample was spotted onto a glow-discharged carbon film 300 mesh copper grid (Electron Microscopy Sciences), allowed to incubate for 2 min, and gently blotted onto a filter paper to remove the solution. The grid was subsequently stained with 5 µL of 2% uranyl acetate for 2 minutes, followed by blotting to remove excess stain. The specimens were analyzed using a FEI Tecnai 12 electron microscope at an accelerating voltage of 120 kV.

### Cryo-EM sample preparation, screening, data collection and processing

Quantifoil Cu R1.2/1.3, 300 mesh were glow-discharged using a Pelco EasiGlow at 15-30 mA for 30-45 seconds. Three-microliter aliquots of extracted fibrils were applied to the grids, blotted with Whatman filter paper for 3-4 seconds (blot force of −1) at 100% humidity, 22°C, and plunged frozen into liquid ethane using a Thermo Fisher Vitrobot Mark IV. Cryo-EM samples were screened using either the Talos Arctica or Glacios at the Cryo-Electron Microscopy Facility (CEMF) at University of Texas Southwestern Medical Center (UTSW). Datasets were collected on a 300 kV Titan Krios microscope (FEI) at Stanford-SLAC Cryo-EM Center (S2C2) using a Falcon 4i detector (Supplementary Table 1). Details on pixel size, frame rate, dose rate, final dose, and number of micrographs per sample are provided in Supplementary Table 1. Automated data collection was performed by EPU3.5 imaging software (Thermo Fisher Scientific).

The raw movie frames were gain-corrected, motion-corrected and dose-weighted using RELION’s own implemented motion correction program^26^. Contrast transfer function (CTF) estimation was carried out using CTFFIND 4.1^27^. All steps of helical reconstruction, three-dimensional (3D) refinement, and post-process were carried out using RELION 4.0^28^.

For both datasets, filaments were initially manually picked from a small subset of 100 movies, resulting in the extraction of approximately 7000-10000 segments. These segments were used to train the neural network Topaz for autopicking^29^. Topaz picked particles were extracted with box sizes of 1024 or 256 pixels, then downscaled to 256 or 128 pixels, respectively. Helical parameters were estimated using reference-free 2D classification of a larger box size, while 2D classification using a smaller box size was used to select particles for further processing. The fibril helix was assumed to be left-handed. The initial 3D classification was performed with an average of ~30k to 40k particles per class using an elongated Gaussian blob as a reference. Particles potentially leading to the best reconstructed map were selected for subsequential 3D classifications and 3D auto-refinements. CTF refinements were performed to achieve higher resolution. The final maps were post-processed using recommended standard procedures in RELION^28^. The overall resolution estimate was evaluated based on the FSC at a 0.143 threshold between two independently refined half-maps.

## Model building

A previously published model of ATTRv-V30M (pdb code 6SDZ) was used as the template to build all near atomic resolution models. Rigid body fit zone, and real space refine zone were performed to obtain the resulting models using COOT^30^. Further refinement was carried out using phenix.real_space_refine from PHENIX 1.20. All refinement statistics are summarized in Supplementary Table 1.

## Stabilization energy calculation

The stabilization energy per residue was calculated by the sum of the products of the area buried for each atom and the corresponding atomic solvation parameters^18^ (Supplementary Figure 4). The overall energy was calculated by the sum of energies of all residues, and assorted colors were assigned to each residue, instead of each atom, in the solvation energy map.

**Figure Panels**. All figure panels were created with Adobe Illustrator.

## Acknowledgments

In memory of Dr. Merrill D. Benson, whose significant contributions have advanced our understanding of amyloid diseases and provided invaluable support to affected families for many years. We are grateful to the patients and families who generously donated tissues and to the University of Indiana for providing these materials. Our thanks also go to Dr. Michael Sawaya for his invaluable advice. We appreciate the engaging discussions and insightful feedback from SBWIP and UTSW. We are thankful to the UTSW Cryo-Electron Microscopy Facility, UTSW Structural Biology Laboratory, and UTSW Electron Microscopy Core Facility, as well as the national cryo-EM facilities at Stanford-SLAC (project CA60) and PNCC (project 51267) for their instrumental and technical support, and data collection efforts. We also extend our gratitude to the UTSW Proteomics core for their technical support in the proteomics experiments.

## Funding

American Heart Association, Career Development Award 847236, L.S.

National Institutes of Health, National Heart, Lung, and Blood Institute, New Innovator Award DP2-HL163810, L.S.

Welch Foundation, Research Award I-2121-20220331, L.S.

UTSW Endowment, Distinguished Researcher Award from President’s Research Council and start-up funds, L.S.

Cryo-EM research was partially supported by the following grants:

National Institutes of Health grant U24GM129547, Department of Energy Office of Science User Facility sponsored by the Office of Biological and Environmental Research

Department of Energy, Laboratory Directed Research and Development program at SLAC National Accelerator Laboratory, under contract DE-AC02-76SF00515

NIH Common Fund Transformative High Resolution Cryo-Electron Microscopy program (U24 GM129539)

The Cryo-Electron Microscopy Facility and the Structural Biology Laboratory at UTSW are supported by a grant from the Cancer Prevention & Research Institute of Texas (RP170644).

The Electron Microscopy Core Facility at UTSW is supported by the National Institutes of Health (NIH) (1S10OD021685-01A1 and 1S10OD020103-01).

Part of the computational resources were provided by the BioHPC supercomputing facility located in the Lyda Hill Department of Bioinformatics at UTSW. URL: https://portal.biohpc.swmed.edu.

## Author contributions

Conceptualization: B.N., L.S. S.A. Y.A

Methodology: B.N., S.A., A.Y., L.S.

Investigation: B.N., S.A., A.Y., V.S., J.V.A., P.K., P.S., M.P., Y.A., M.C.F.R., R.P., P.B., L.W., A.L., L.V., B.K.B., L.S

Visualization: B.N., S.A., L.S.

Funding acquisition: L.S.

Project administration L.S.

Supervision: B.N., S.A., L.S.

Writing – original draft: S.A, B.N

Writing – review & editing: B.N., S.A., A.Y., L.S.

## Data and Materials Availability

Structural data have been deposited into the Worldwide Protein Data Bank (wwPDB) and the Electron Microscopy Data Bank (EMDB) with the following EMD accession codes: To be released (ATTRv-V30M-heart), To be released (ATTRv-V30M-nerve), and PDB accession codes: To be released (ATTRv-V30M-heart), To be released (ATTRv-V30M-nerve). The PDB accession code for the previously reported coordinates of ATTRv-V30M used for data processing is 6SDZ. All data generated or analyzed during this study that support the findings are available within this published article and its supplementary data files. Graphed data is provided in the source data file. Cardiac and nerve specimens were obtained from the laboratory of the late Dr. Merrill D. Benson at Indiana University. These specimens are under a material transfer agreement with Indiana University and cannot be distributed freely.

