## Supplemental Figures and Table for "ATTRv-V30M Type A amyloid fibrils from heart and nerves exhibit structural homogeneity"

**Supplementary figures:**

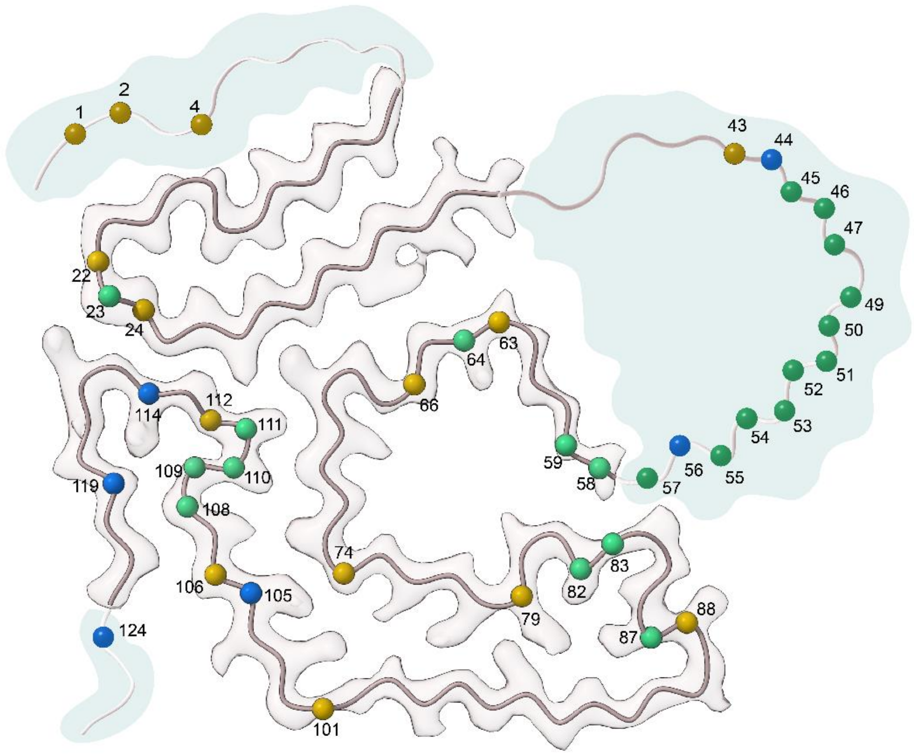


**Supplementary Fig 1. Schematic illustration of potential proteolytic sites determined from MS analysis of non-tryptic peptides**. Blue dots represent cleavage sites unique to ATTRv-V30M nerve fibrils, orange represents cleavage sites unique to ATTRv-V30M heart fibrils and green represents common cleavage sites found in both the cardiac and nerve fibrils.

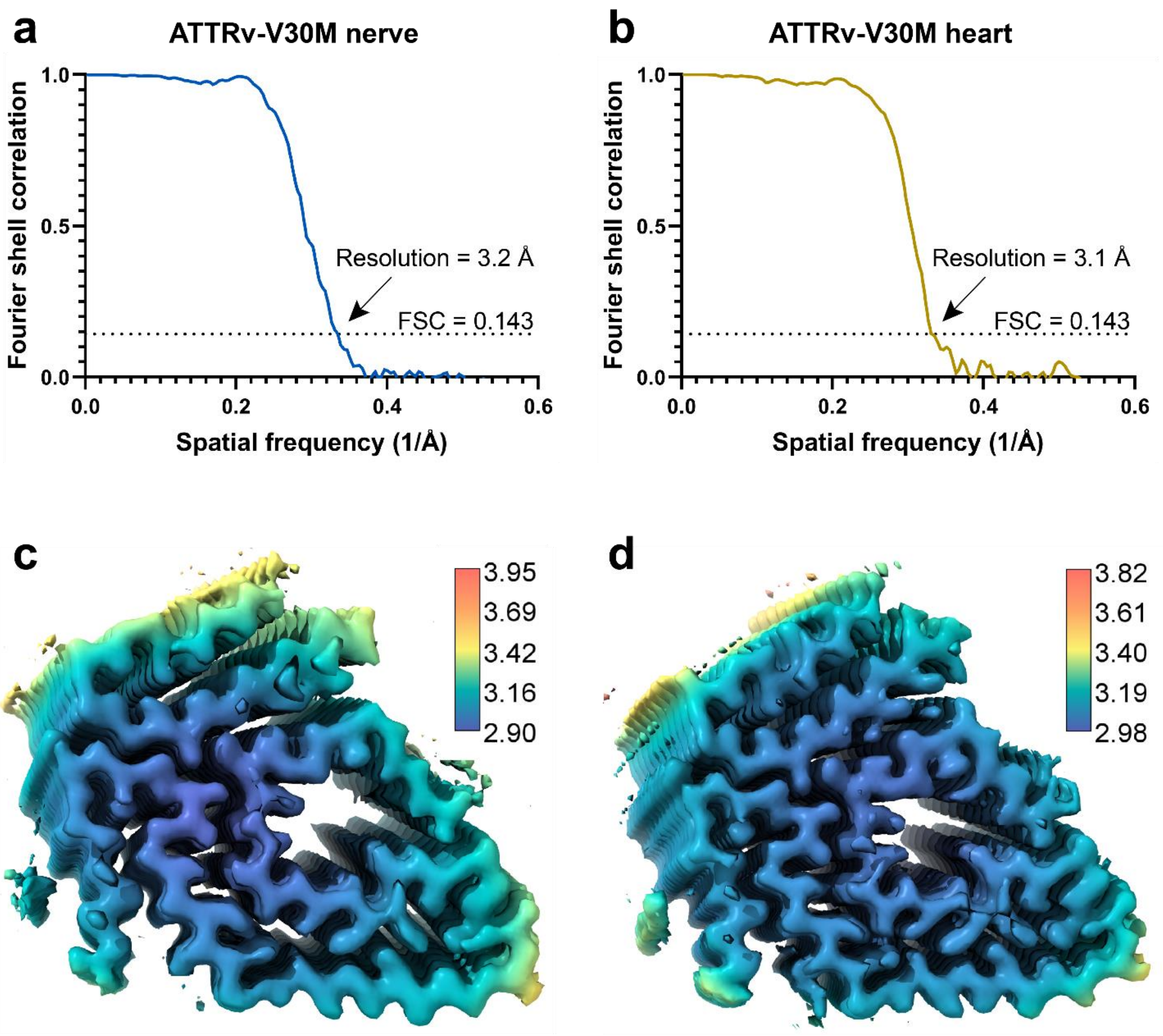


**Supplementary Fig 2: Evaluation of resolution of cryo-EM maps**. FSC curves derived from two independently analyzed half maps for ATTRv-V30M (a) nerve and (b) heart fibril structure. Local resolution estimation for 3D reconstructions of ATTR fibrils, measured in angstroms, for (c) nerve and (d) heart fibril structure.

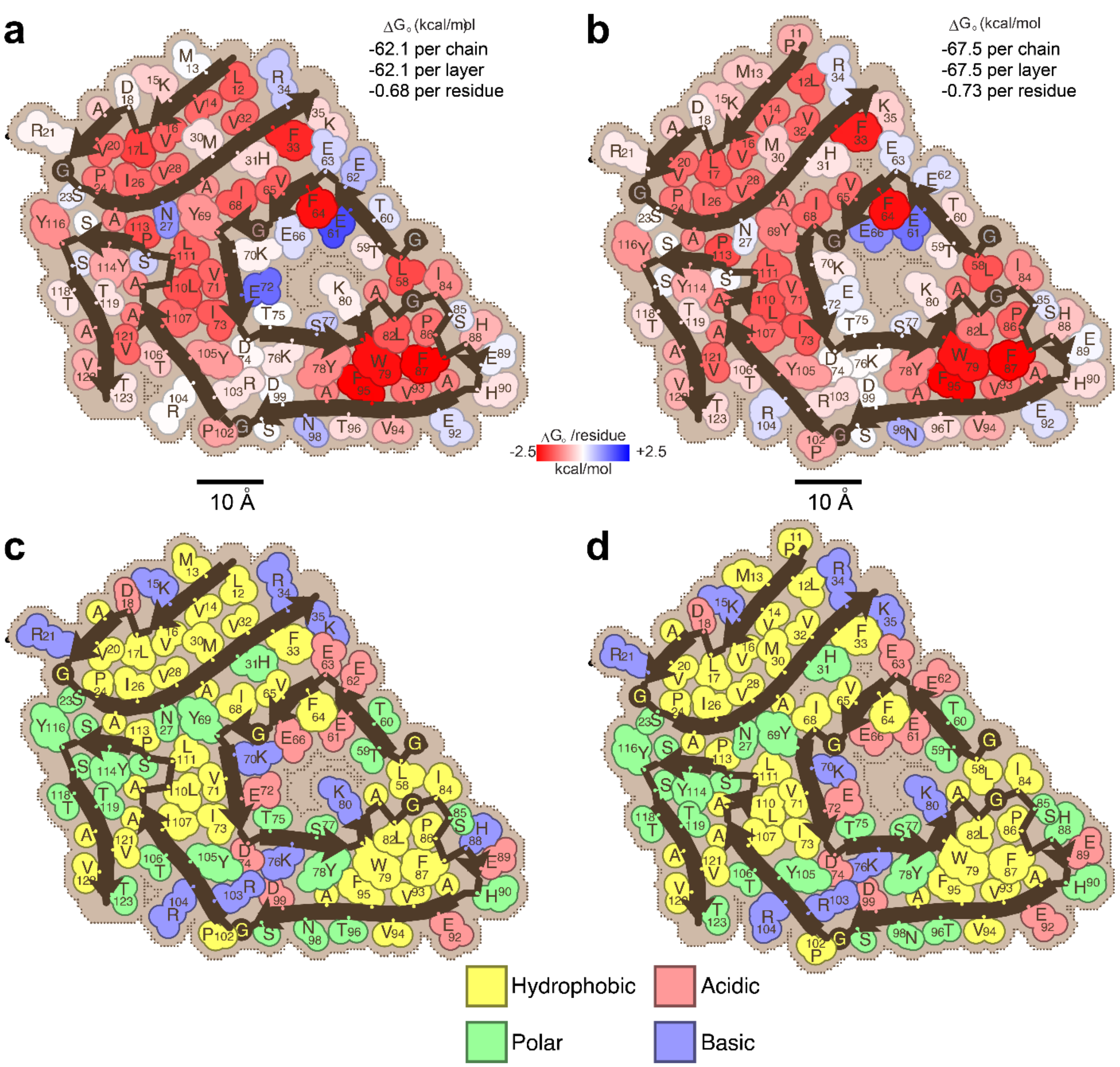


**Supplementary Fig 3: Solvation energy and residue composition analysis of nerve and heart ATTR fibril structures.** Representation of solvation energies per residue for fibril structure from (a) nerve and (b) heart. Residues depicted in red indicate more favorable solvation energies (-2.5 kcal/mol), whereas residues in blue represent less favorable stabilization energies (2.5 kcal/mol). Scale: 10Å. Schematic diagram of ATTRv-V30M fibril structures displaying the composition of residues in (c) nerve and (d) heart. The residues are color-coded based on amino acid categories, as described in the labels.

**Supplementary Table 1.** Data collection and model refinement statistics.

| Data collection of ATTRv-V30M | Nerve | Heart |
|---|---|---|
| **Microscope (Model)** | Titan Krios (G3i) | Titan Krios (G3i) |
| **Acceleration Voltage (kV)** | 300 | 300 |
| **Detector** | Falcon 4i | Falcon 4i |
| **Software** | EPU 3.5 | EPU 3.5 |
| **Magnification** | 130,000 | 130,000 |
| **Pixel size at detector (Å/px)** | 0.946 | 0.946 |
| **Defocus range (µm)** | 0.9 to 2.1 | 0.9 to 2.1 |
| **Total dose (e)** | 40 | 40 |
| **Exposure time (sec)** | 4.57 | 4.98 |
| **Number of movie frames** | 40 | 40 |
| **Usable micrographs** | 8,957 | 5751 |
| **Box size** | 256 | 256 |
| **Total extracted segments** | 1,064,689 | 2,311,260 |
| **Number of segments after 2D** | 774,031 | 539,031 |
| **Number of straight segments** | 16,615 | 40,533 |
| **Number of segments after 3D** | 104,108 | 99,731 |
| **Symmetry imposed** | C1 | C1 |
| **Helical rise (Å)** | 4.93 | 4.90 |
| **Helical twist (°)** | -1.31 | -1.23 |
| **Crossover length (Å)** | 677 | 717 |
| **B factor** | -118.4 | -109.4 |
| **Map resolution (Å; FSC=0.143)** | 3.36 | 3.14 |
| **Map resolution (Å; FSC=0.5)** | 3.7 | 3.6 |
| **Non-hydrogen atoms** | 3560 | 3595 |
| **Protein residues** | 455 | 460 |
| **Number of chains** | 5 | 5 |
| **Water/ligands** | 0 | 0 |
| **MolProbity score** | 1.86 | 1.75 |
| **Clash score** | 10.41 | 7.93 |
| **Rotamer outliers (%)** | 0 | 0 |
| **R.M.S deviations bonds (Å)** | 0.006 | 0.003 |
| **R.M.S deviations angle (°)** | 0.658 | 0.762 |
| **Ramachandran plot** (5) | | |
| Favored | 95.40 | 95.45 |
| Allowed | 4.60 | 4.55 |

| | | |
|---|---|---|
| Outliers | 0.00 | 0.00 |
| **CaBLAM outliers (%)** | 4.82 | 3.57 |
| **Model vs Data (%)** | 0.84 | 0.86 |
